# Integrative analysis of multi-platform reverse-phase protein array data for the pharmacodynamic assessment of response to targeted therapies

**DOI:** 10.1101/769158

**Authors:** Adam Byron, Stephan Bernhardt, Bérèngere Ouine, Aurélie Cartier, Kenneth G. Macleod, Neil O. Carragher, Vonick Sibut, Ulrike Korf, Bryan Serrels, Leanne de Koning

## Abstract

Reverse-phase protein array (RPPA) technology uses panels of high-specificity antibodies to measure proteins and protein post-translational modifications in cells and tissues. The approach offers sensitive and precise quantification of large numbers of samples and has thus found applications in the analysis of clinical and pre-clinical samples. For effective integration into drug development and clinical practice, robust assays with consistent results are essential. Leveraging a collaborative RPPA model, we set out to assess the variability between three different RPPA platforms using distinct instrument set-ups and workflows. Employing multiple RPPA-based approaches operated across distinct laboratories, we characterised a range of human breast cancer cells and their protein-level responses to two clinically relevant cancer drugs. We integrated multi-platform RPPA data and used unsupervised learning to identify protein expression and phosphorylation signatures that were not dependent on RPPA platform and analysis workflow. Our findings indicate that proteomic analyses of cancer cell lines using different RPPA platforms can identify concordant profiles of response to pharmacological inhibition, including when using different antibodies to measure the same target antigens. These results highlight the robustness and the reproducibility of RPPA technology and its capacity to identify protein markers of disease or response to therapy.

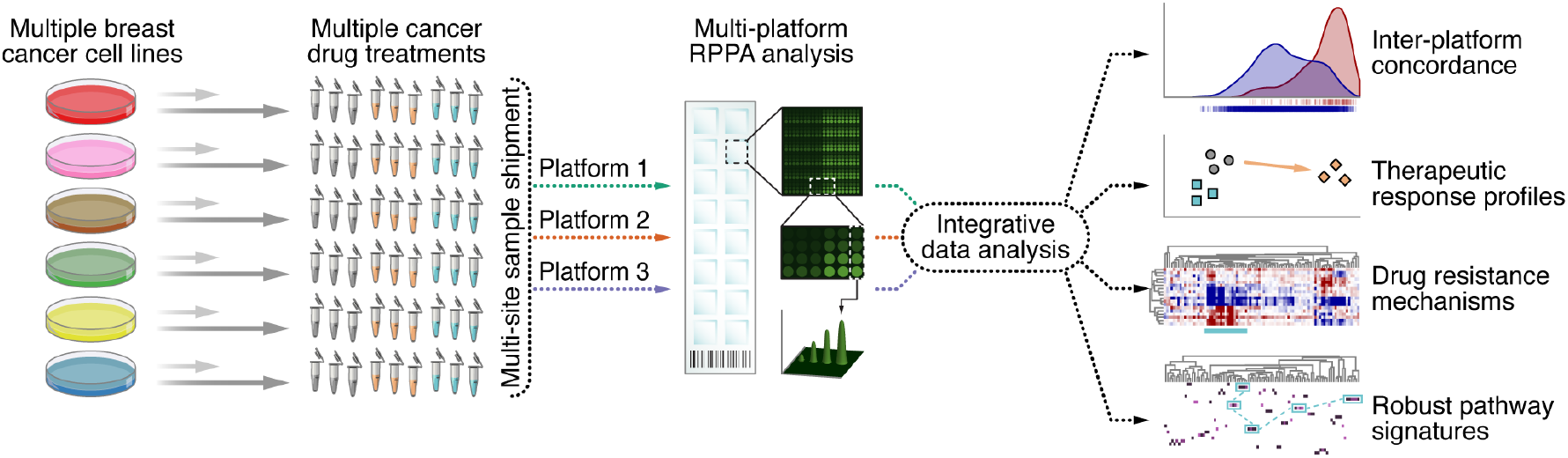

## Introduction

In the era of personalised medicine and targeted cancer therapies, identifying those patients that will benefit from existing and new therapies is paramount. Genetics is already being used to assist clinical decision-making in specific cases (Aronson & Rehm 2015), but additional levels of biological information are required to better understand disease and more accurately predict phenotype from genotype (Friedman et al. 2015). A crucial source of information in this context is the proteome and, notably, the activation status of dynamic cell signalling pathways through post-translational protein modifications. Indeed, genomic mutations are not always associated with activated signalling pathways and, conversely, pathway activation can occur in the absence of mutations, as exemplified by *PIK3CA* (Cancer Genome Atlas Network 2012) and human epidermal growth factor receptor 2 (Her2, also known as ErbB2) signalling (Wulfkuhle et al. 2012) in breast cancer.

Several technologies exist for protein biomarker discovery and validation (Mueller et al. 2018, Giudice & Petsalaki 2019, Pierobon et al. 2019), among which reverse-phase protein array (RPPA) is a technology of choice for its unequalled sample throughput. The RPPA technology uses panels of monospecific affinity reagents (usually validated, high-quality antibodies) to quantify, with high precision and sensitivity, the abundance of specific proteins and their post-translationally modified forms in biological specimens (Paweletz et al. 2001, Akbani et al. 2014). Protein samples derived from cells or tissues are immobilised on a solid substrate, deposited as small spots on multiple arrays, and each array is probed with a single, epitope-specific antibody. This enables simultaneous quantification of multiple proteins and post-translational modifications in hundreds of samples, a multiplex capability not available in any other current proteomic technology. The capacity to analyse large sample numbers enables analysis of multiple sample conditions, such as drug treatments, dose responses and time courses, resulting in data series that can support systems biology and drug discovery pipelines (Macleod et al. 2017, Hsieh et al. 2018). The high sensitivity (in the picomole–femtomole range) and good reproducibility of RPPA technology (Paweletz et al. 2001, Ramaswamy et al. 2005, Tibes et al. 2006, Grote et al. 2008, Dupuy et al. 2009, Troncale et al. 2012) have motivated its application to a wide range of sample types, including cell lines, preclinical (e.g. xenograft) models and patient-derived material. Indeed, the microscale printing of very small amounts of samples is of particular benefit for analysis of limited clinical or preclinical material, and RPPA has become a powerful addition to the biomedical analytical toolbox for the investigation of disease mechanisms, diagnostics and prognostics, notably in cancer (Grubb et al. 2009, Gonzalez-Angulo et al. 2011, Murakoshi et al. 2011, Hayashi et al. 2014, Bernhardt et al. 2017, Hutter et al. 2017, Lièvre et al. 2017, Aslan et al. 2018, Faham et al. 2018, Teo et al. 2018).

The RPPA workflow is composed of several distinct steps, which can each be adapted to the needs of the laboratory or the study. RPPAs thus offer a highly flexible, modular proteomic technology, enabling numerous possible technical set-ups and protocols. As most laboratories using RPPA technology have developed a customised set-up, there are essentially as many workflows as there are RPPA platforms. Differences between platforms are diverse and can include the type of printer used to create the arrays, sample spotting conditions, slide substrate chemistry, primary and secondary antibodies used for immunostaining and slide scanner optics (Byron 2019). Moreover, no standard tools exist to quantify, normalise and quality control RPPA data. Several RPPA data processing methods have been developed (Mircean et al. 2005, Hu et al. 2007, Anderson et al. 2009, Neeley et al. 2009, Zhang et al. 2009, Mannsperger et al. 2010, Li et al. 2012, Neeley et al. 2012, Troncale et al. 2012, Kaushik et al. 2014, List et al. 2014, Liu et al. 2014, Ju et al. 2015, Sun et al. 2015), but these are often designed for specific technical set-ups, requiring, for example, particular array layouts or raw data formats. To our knowledge, no extensive cross-platform validation has been reported for RPPA technology. There is, therefore, a need to assess and understand the impact of variation between RPPA data derived from different platforms across international research centres.

As RPPA workflows are being developed for use in clinical settings, for which robust assays are paramount (Gallagher & Espina 2014, Masuda & Yamada 2015), there is a growing need for the evaluation of the reproducibility of RPPA platform outputs. Herein, we use collaborative RPPA-based proteomics, employing distinct RPPA workflows at multiple research sites in different countries, to characterise a range of human breast cancer cell lines and their biochemical responses to two clinically relevant cancer drugs. We combine data derived from different RPPA platforms to assess inter-platform variation and use integrative analysis to identify platform-independent protein markers of response to drug treatment. Such cross-platform validation may thus have utility in multi-centre evaluation of robust protein markers of disease and therapeutic response.

## Results

### Multi-platform RPPA analysis of breast cancer cell lines

To assess the reproducibility of RPPA technology across multiple laboratories, we developed an international multi-platform approach that integrated RPPA data derived from three research sites across Europe (Paris, France; Heidelberg, Germany; Edinburgh, United Kingdom). We selected for this study six breast cancer cell lines, encompassing different breast cancer molecular subtypes and presenting distinct drug sensitivities (Supplementary Table 1). The cells were cultured in the absence or presence of two kinase inhibitors for 20 minutes or 24 hours and lysed in biological triplicate. In total, we generated 108 snap-frozen lysates, which were shipped to the three research sites (Fig. 1a). Samples were analysed at each site using the respective in-house RPPA platforms, the set-up of which differed at many stages of the RPPA analysis workflow, including slide type, the number of technical replicates and dilutions per sample, read-out dye, scanner, image analysis software and normalisation procedure (Supplementary Table 2), enabling the capture of variation between RPPA platforms operated in different laboratories. Microarrayed samples were probed with panels of validated antibodies in routine use on the three RPPA platforms. To enable dataset comparison, all antibodies were assigned unique antibody identifiers, and only data derived from antibodies targeting the same protein(s) or phosphorylated residue(s) (including different antibodies from different suppliers) acquired on all three RPPA platforms were used for further analysis (Supplementary Table 3). This experimental design thus enabled the assessment of inter-platform concordance.

**Figure 1.**
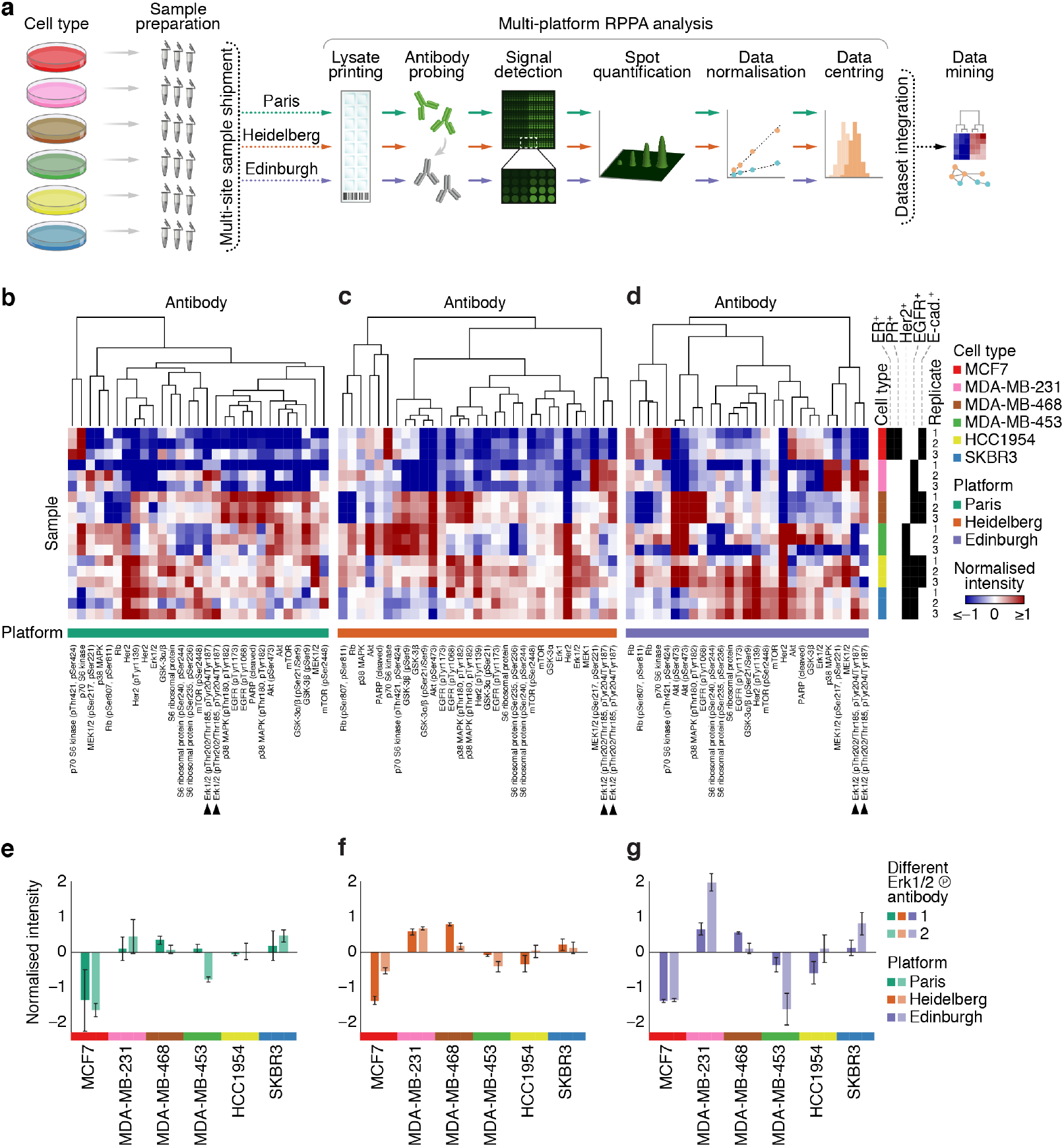
Multi-platform RPPA analysis of breast cancer cell lines. (**a**) Schematic illustration of the multi-platform RPPA workflow. (**b**–**d**) Hierarchical cluster analysis of RPPA data of six breast cancer cell lines cultured under control conditions derived from three RPPA platforms. RPPA data derived from the Paris (**b**), Heidelberg (**c**) and Edinburgh (**d**) platforms were clustered on the basis of antibody-wise dissimilarity (Spearman rank correlation coefficient– based distance). Clustering was performed separately for each platform, and sample order was maintained for each cluster analysis. Annotation bars indicate cell type, cell type receptor status and RPPA platform. Arrowheads indicate different phosphorylated Erk1/2 antibodies. (**e**–**g**) RPPA data for phosphorylated Erk1/2 (pThr202/Thr185, pTyr204/Tyr187) derived from the Paris (**e**), Heidelberg (**f**) and Edinburgh (**g**) RPPA platforms. All three platforms used the same two antibodies that recognise phosphorylated Erk1/2 (dark bars, antibody identifier Erk1/2_pThr202/Thr185,pTyr204/Tyr187_a; light bars, antibody identifier Erk1/2_pThr202/Thr185,pTyr204/Tyr187_b). Annotation bars (*x*-axis) indicate cell type. Data are means ± s.e.m. (*n* = 3 independent samples). For further details, see Supplementary Fig. 1 and Supplementary Table 4.

First, RPPA data of the six breast cancer cell lines cultured under control conditions were analysed using the respective in-house data analysis procedures of the three research sites. Analysis of these control samples generated in biological triplicate with antibodies used in the multi-platform RPPA analysis resulted in 522, 558 and 486 antibody readings (processed signal intensities) for the Paris, Heidelberg and Edinburgh platforms, respectively. Normalised RPPA data derived from each platform were then clustered antibody-wise to identify similarities between validated antibodies (Supplementary Table 4). For each RPPA platform, clusters containing antibodies targeting the same protein or phosphoprotein were identified, indicating that normalised intensity profiles across cell types correlated for subsets of different antibodies (Fig. 1b–d, Supplementary Fig. 1). Most distinct antibodies targeting the same antigen, such as phosphorylated Erk1/2, clustered together in each RPPA dataset, indicating consistent results for the same antigen (Fig. 1e–g, Supplementary Fig. 1). In addition, antibodies targeting different phosphorylated residues on the same protein or antibodies targeting phosphorylated and corresponding total proteins, such as for S6 ribosomal protein or mTOR, generally clustered together in each RPPA dataset, although degree of correlation varied between RPPA platforms and distance metrics (Fig. 1b–d, Supplementary Fig. 1). These data suggest that many of the antibodies routinely used for RPPA analysis at the three research sites in this study give comparable results relative to the rest of each dataset.

### Integrative analysis of multi-platform RPPA data

To assess the comparability of RPPA data originating from multiple platforms, we integrated normalised RPPA data for control samples derived from each platform (1,566 antibody readings) and analysed the integrated dataset using unsupervised learning. For dataset integration, antigens targeted by each antibody were classified, capturing antibody recognition of related protein isoforms or family members where applicable (e.g. antibodies recognising Erk1 and Erk1/2 were linked and assigned the same antibody antigen class, Erk1/2). Each data point was linked to the RPPA platform from which it was derived to enable downstream data analysis. We reduced the dimensionality of the integrated multi-platform RPPA dataset using principal component analysis. This unsupervised analysis of the six breast cancer cell lines revealed separation in feature space for several of the cell types, indicating cell type–specific expression of the proteins and phosphoproteins measured (Fig. 2a). Next, two-dimensional hierarchical cluster analysis of the integrated dataset, using multiple distinct distance functions to quantify dissimilarity between data points in the feature space, partitioned samples by breast cancer cell line, elucidating discriminatory profiles of protein and phosphoprotein expression for each cell type (Fig. 2b, Supplementary Fig. 2). Moreover, unsupervised cluster analysis identified correlated subsets of antibodies targeting the same antigens. The data driving these antibody clusters were generally derived from multiple RPPA platforms and, where available, using different antibodies (Fig. 2b, Supplementary Table 5), suggesting that the profiles of protein and phosphoprotein expression were comparable between RPPA platform set-ups.

**Figure 2.**
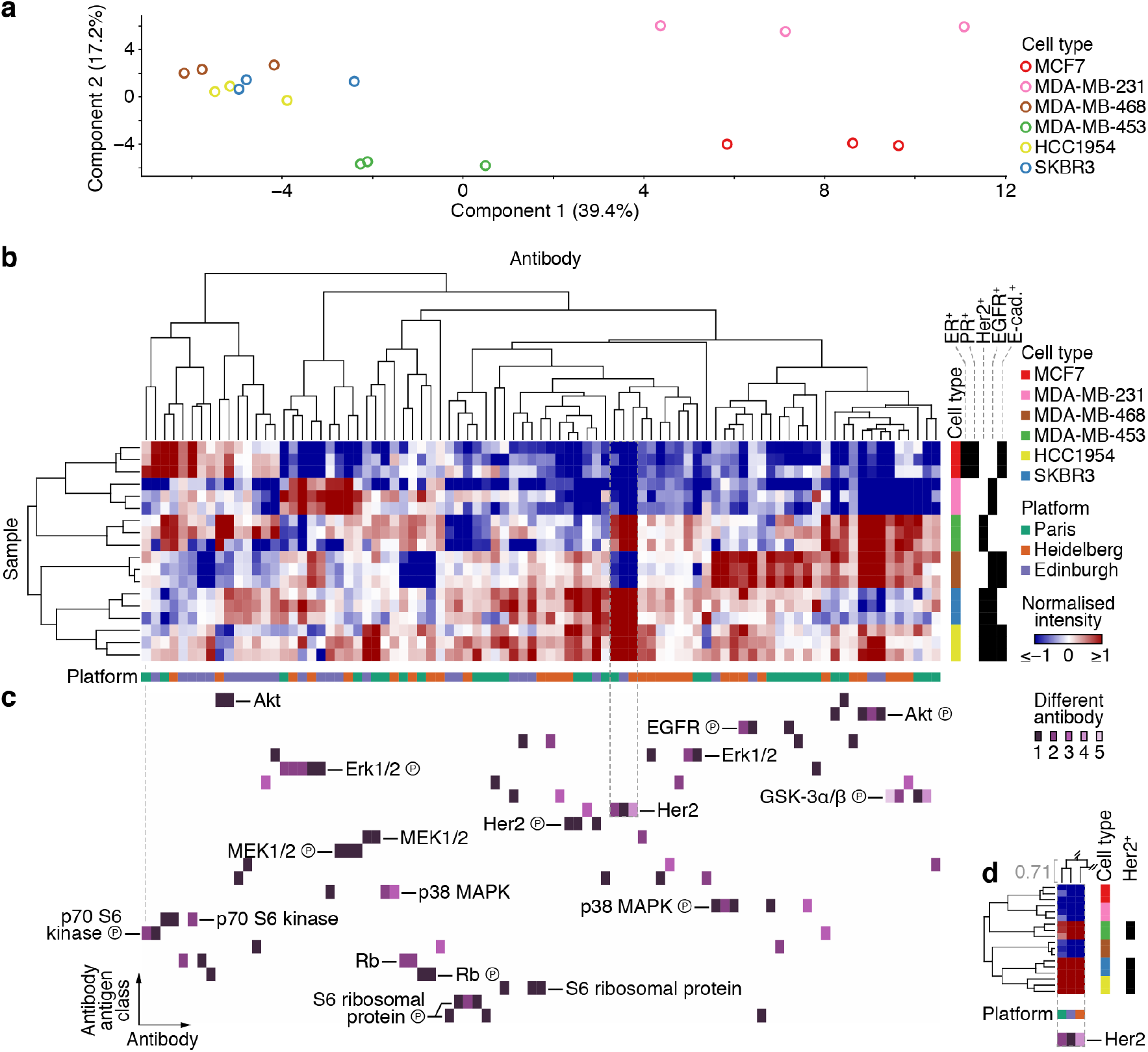
Integrative analysis of multi-platform RPPA data. (**a**) Principal component analysis of integrated multi-platform RPPA data of six breast cancer cell lines cultured under control conditions. (**b**) Integrated multi-platform RPPA data were clustered on the basis of antibody-wise and sample-wise dissimilarities (Spearman rank correlation coefficient–based distance). Annotation bars indicate cell type, cell type receptor status and RPPA platform. (**c**) Clustered antibody antigen map for all antibodies used in the integrative analysis. The map is aligned with the hierarchical clustering results in **b** (grey dashed line indicates alignment of first antibody (column) end-node). Distinct antibodies that target the same antigen class (unique antibody identifiers) are indicated by different shades of purple. Antibody antigen classes are ordered alphabetically for clarity. (**d**) Exemplar of clustered antibodies that recognise Her2, indicated by grey dashed box in **b** and **c**. Antibodies used by all three RPPA platforms clustered in antibody– antibody antigen space. Spearman rank correlation coefficient of the cluster column node is shown in grey. All three platforms used different antibodies that recognise Her2 (antibody identifiers for Paris, Her2_b; Heidelberg, Her2_d; Edinburgh, Her2_a). For further details, see Supplementary Fig. 2 and Supplementary Table 5.

To interrogate the similarity between antigen expression profiles for distinct antibodies used on different RPPA platforms, we devised a data-driven representation of antibody similarity, which we termed a clustered antibody antigen map. Antibodies used at each RPPA platform (clustered as for Fig. 2b) were annotated with their respective antigens (classified as described in Methods). With clustered antibodies as columns, antigen annotations were expanded into rows of a matrix. The resulting matrix was populated with unique antibody identifiers to distinguish different antibodies, from which a heatmap of clustered antibody–antigen space was generated. Thus, the antibody antigen map provides detailed annotation of cluster analysis of multi-platform RPPA data. This enables identification of clustered antibodies that recognise antigens with similar expression profiles across the integrated dataset (Fig. 2b,c, Supplementary Fig. 2).

The antibody antigen map revealed that the expression of many of the antigens tested (71%) clustered with expression of the same antigen determined by at least one other RPPA platform (Fig. 2c,d). In addition, the expression of several of the antigens for which distinct antibodies were used (61%) clustered with the same antigen detected by a different antibody (Fig. 2c,d). Together, these results indicate that RPPA analyses performed at different research sites using distinct setups can identify concordant sets of distinct antibodies that target the same antigens. This implies that different high-quality, validated antibodies can be used to generate consistent results from the same samples using different RPPA platforms.

### Consistency of multi-platform RPPA analysis of drug-treated breast cancer cell lines

To analyse the robustness of RPPA technology using a more relevant ‘intervention’ dataset (i.e. including treatment conditions), we extended the integrative analysis to include multi-platform RPPA data of the six breast cancer cell lines treated with two clinically relevant drugs, lapatinib (Tykerb, Tyverb) and selumetinib (AZD6244, ARRY-142886). Lapatinib is a reversible ATP-mimetic tyrosine kinase inhibitor of epidermal growth factor receptor (EGFR, also known as ErbB1) and Her2 (Spector et al. 2005); selumetinib is a selective ATP-independent allosteric inhibitor of mitogen-activated protein kinase kinase (MAPKK, also known as MEK) (Yeh et al. 2007). To capture the signalling dynamics of the proteins under investigation, cells were treated with either drug or with DMSO (vehicle control) for 20 min and 24 h (Fig. 3a).

**Figure 3.**
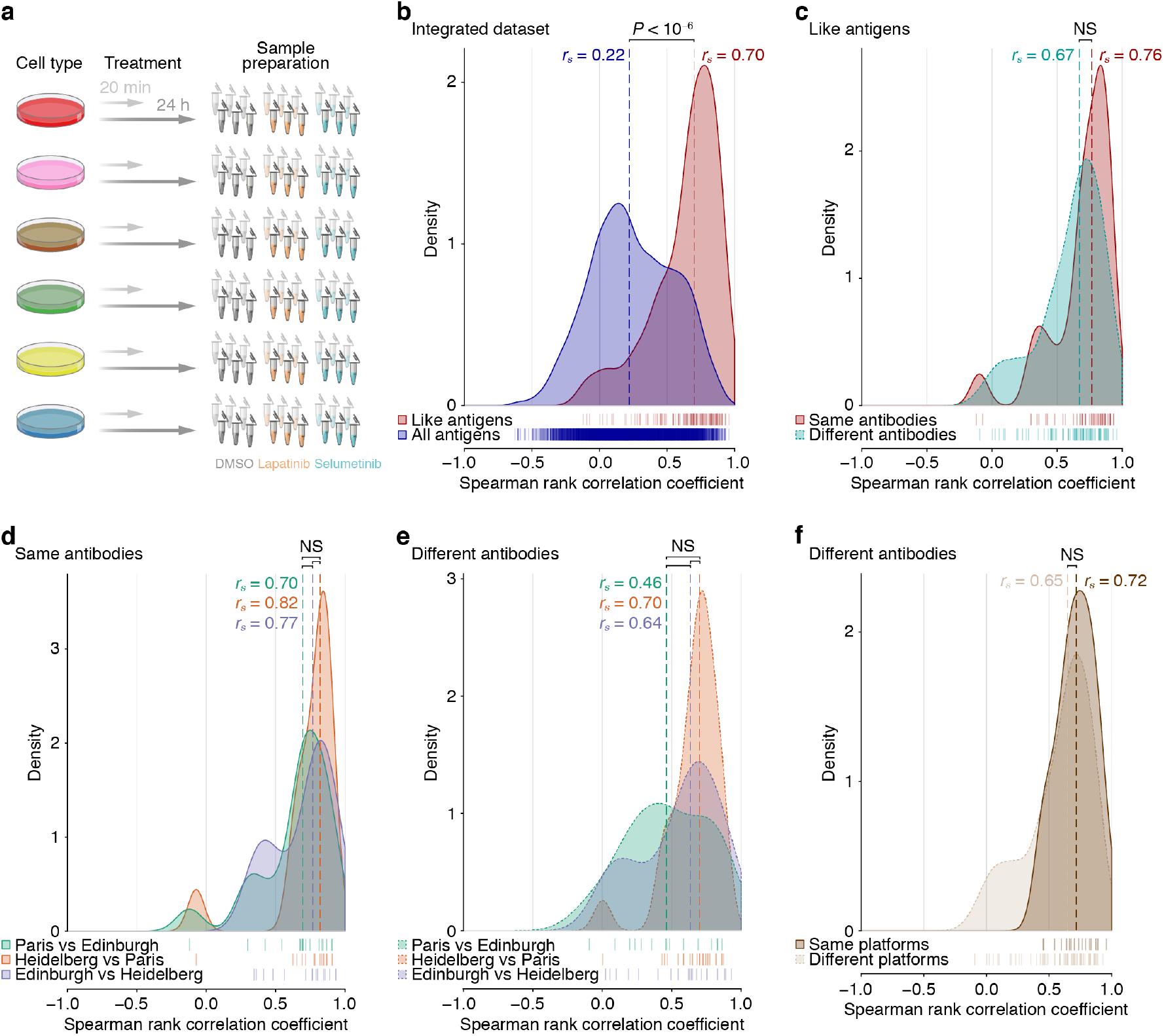
Correlations of RPPA data used for integrative analysis of drug-treated breast cancer cell lines. (**a**) Schematic illustration of the drug treatment experiment. Samples were processed as part of the same workflow shown in Fig. 1. (**b**) Correlations of RPPA data derived from antibodies recognising the same antigen class (like antigens) were compared to those derived from all antibodies used in the integrated multi-platform RPPA dataset. (**c**) For like antigens, correlations of RPPA data derived from the same antibodies were compared to those derived from different antibodies. (**d**,**e**) Correlations between RPPA data generated at the different RPPA platforms were compared for data derived from the same antibodies (**d**) and different antibodies recognising like antigens (**e**). (**f**) Correlations of RPPA data derived from different antibodies recognising like antigens generated on the same RPPA platform were compared to those generated on different RPPA platforms. For **b**–**f**, kernel density estimates of Spearman rank correlation coefficients for every pair-wise combination of unique antibody identifiers were computed. Spearman rank correlation coefficient data points for each set of comparisons are indicated by rug plots. For each set of comparisons, the median Spearman rank correlation coefficient (*r*_*s*_) is shown (dashed lines). NS, not significant. For further details, see Supplementary Fig. 3.

First, to examine the consistency of results generated by the antibodies used in the multi-platform RPPA analysis, we calculated correlations between all-sample RPPA data derived from all antibodies tested, which consisted of 9,396 antibody readings. This analysis showed that RPPA data derived from antibodies recognising the same antigen class (i.e. ‘like’ antigens) were generally well correlated (median Spearman rank correlation coefficient, *r*_*s*_ = 0.70) (Fig. 3b). In contrast, data derived from all antibodies – regardless of target – were generally poorly correlated (*r*_*s*_ = 0.22), as expected (Fig. 3b), implying that RPPA-based quantification of like target antigens is in substantially better agreement than quantification of random antigens in the dataset. Notably, RPPA data for antigens recognised by the same antibody were correlated to a similar level to those recognised by different antibodies (Fig. 3c), indicating that distinct, validated antibodies generate consistent results from the same samples. In addition, correlations between normalised RPPA data derived from all antibodies were lower than those between corresponding raw RPPA data, resulting in a better separation of correlation distributions for like antigens and for all antibodies (Supplementary Fig. 3). This suggests that normalisation of RPPA data better differentiates concordant data (derived from antibodies recognising the same antigen class) from less-concordant data (derived from all antibodies regardless of target).

To assess the reproducibility of RPPA results across different RPPA platforms, we compared correlation distributions for like antigens for each pair-wise combination of platforms. Each platform comparison showed a similar correlation distribution for antigens recognised by the same antibodies (Fig. 3d) and a similar correlation distribution for antigens recognised by different antibodies (Fig. 3e), although different antibodies used at the Paris and Edinburgh platforms were less well correlated. Importantly, antigens detected by different antibodies used at different RPPA platforms were, in general, almost as well correlated as those used at the same RPPA platform (Fig. 3f). These data show that RPPA analyses of the same samples at different platforms using distinct workflows yield consistent results, including when several different antibodies are used to recognise the same antigen (protein or phosphoprotein) of interest.

### Integrative multi-platform RPPA analysis of drug-treated breast cancer cell lines

We hypothesised that the observed consistency of multi-platform RPPA data would allow robust detection of potential markers of cellular response to signalling pathway inhibition. To confirm overall changes in RPPA data upon drug treatment of breast cancer cells, we reduced the dimensionality of the integrated dataset using principal component analysis. Unsupervised analysis of all cell lines identified shifts in feature space away from control conditions for some drug-treated cells, suggesting cell type–specific differential regulation of proteins and phosphoproteins (Supplementary Fig. 4). For example, the Her2-amplified SKBR3 cell line is highly sensitive to lapatinib (Hegde et al. 2007, Imami et al. 2012), and treatment with lapatinib induced substantial changes in phosphoprotein abundance, including that of phosphorylated Her2 and EGFR and downstream signalling molecules Akt and Erk1/2 (Fig. 4a,b, Supplementary Fig. 4). In contrast, dimensionality-reduced RPPA data for MCF7 cells, which do not overexpress Her2 or EGFR, did not display a large shift in feature space away from control conditions, in keeping with the lack of response to lapatinib treatment of MCF7 cells (Supplementary Fig. 4). For cells treated with selumetinib, a strong reduction in phosphorylated Erk1/2 – which is activated upon phosphorylation by MEK (Dhillon et al. 2007) – was observed in MEK inhibitor–sensitive MDA-MB-231 cells analysed at all RPPA platforms (Fig. 4a,b), whereas SKBR3 cells, which are not as sensitive to selumetinib, displayed a minimal shift in feature space for selumetinib-treated samples (Fig. 4a). In some cell lines, upregulation of phosphorylated MEK1/2 was observed upon selumetinib treatment, particularly after treatment for 24 h, representing the likely effects of reduced negative feedback on the upstream MAPK pathway as a result of transiently inhibited Erk1/2 upon MEK inhibition (Pratilas et al. 2009, Lito et al. 2014). In general, for drug-sensitive cell lines, shifts in feature space were more pronounced for cells treated for 24 h as compared to 20 min, implying that the modulation of signalling pathways was enhanced when cells were challenged with drugs for longer, enabling modelling of the dynamic signalling landscape (Supplementary Fig. 4). Furthermore, in drug-resistant cell lines, we observed the emergence of potential resistance mechanisms, such as the activation of phosphorylated EGFR, Her2 and Akt in SKBR3 cells treated with selumetinib (Supplementary Fig. 4). These analyses show that the RPPA data represent expected changes in breast cancer cell responses to the pharmacological inhibitors tested, capturing relevant signalling dynamics, and serve as a suitable platform for the integrative multi-platform RPPA analysis of drug-treated cells.

**Figure 4.**
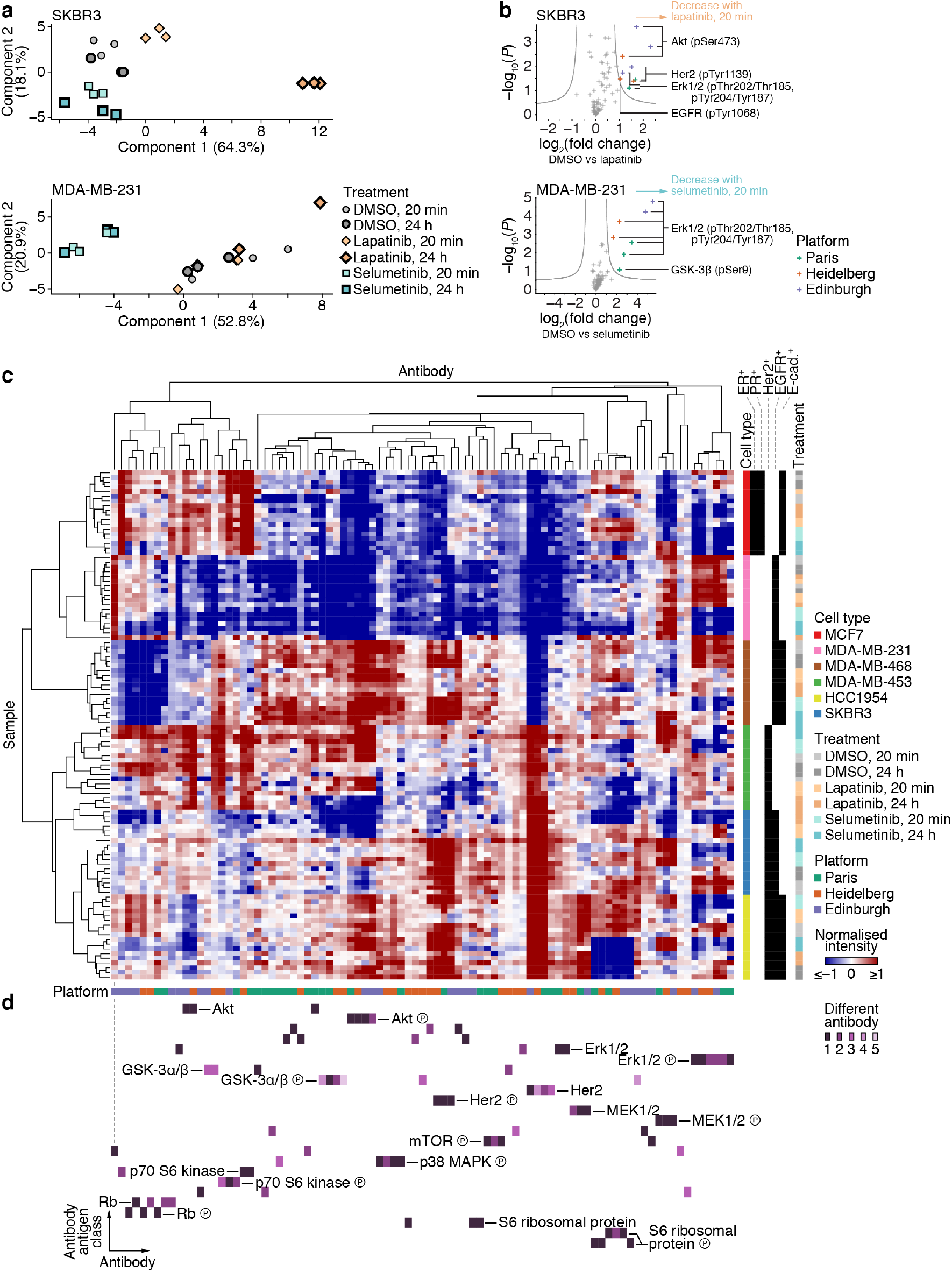
Integrative multi-platform RPPA analysis of drug-treated breast cancer cell lines. (**a**) Principal component analyses of integrated RPPA data of SKBR3 cells (top) or MDA-MB-231 cells (bottom) treated with lapatinib, selumetinib or vehicle control (DMSO) at two timepoints. (**b**) Volcano plots of RPPA data of SKBR3 cells treated with lapatinib (top) or MDA-MB-231 cells treated with selumetinib (bottom) compared to those treated with vehicle control (DMSO) for 20 min. Differentially regulated phosphoproteins are labelled. Grey curves indicate 5% false discovery rate (*n* = 3 independent samples). (**c**) Integrated multi-platform RPPA data of drug-treated breast cancer cell lines were clustered on the basis of antibody-wise and sample-wise dissimilarities (Spearman rank correlation coefficient–based distance). Annotation bars indicate cell type, cell type receptor status, drug treatment and RPPA platform. (**d**) Clustered antibody antigen map for all antibodies used in the integrative analysis. The map is aligned with the hierarchical clustering results in **c** (grey dashed line indicates alignment of first antibody (column) end-node) and coloured and ordered as for Fig. 2. For further details, see Supplementary Figs 4 and 5 and Supplementary Table 6.

We next analysed the integrated RPPA dataset of drug-treated cell lines using unsupervised cluster analysis. Two-dimensional hierarchical clustering partitioned samples by breast cancer cell line, which were further partitioned by treatment condition (for respective drug-sensitive cells), with replicate samples clustering together, indicating robust data-driven grouping of the dataset (Fig. 4c, Supplementary Fig. 5, Supplementary Table 6). Clustered antibody antigen mapping, driven by this unsupervised dataset partitioning, identified clusters of antibodies targeting the same antigen (Fig. 4d, Supplementary Fig. 5). As for the analysis of the cells cultured under control conditions (Fig. 2, Supplementary Fig. 2), expression of many of the antigens tested (75%) clustered with expression of the same antigen determined by at least one other RPPA platform (Fig. 4d). Furthermore, the expression of several of the antigens for which distinct antibodies were used (56%) clustered with the same antigen detected by a different antibody (Fig. 4d). Together, these results indicate that integrative RPPA analysis of drug-treated cells can identify concordant profiles of response to pharmacological inhibition using distinct antibodies and different RPPA platforms. This suggests that the robustness of RPPA technology is suitable for the characterisation of pathway signalling networks across international laboratories.

## Discussion

With RPPA technology poised for adoption into routine clinical laboratory assays, there is a need to assess reproducibility and variation between RPPA data derived from different platforms. Herein, we employed a collaborative RPPA-based proteomics approach to evaluate the consistency of results obtained from the same samples using different RPPA workflows. We combined the RPPA data derived from multiple research sites to assess inter-platform variation and found that different RPPA platforms using distinct set-ups yield remarkably consistent results, including when several different antibodies are used to recognise the same antigen (protein or phosphoprotein) of interest. Indeed, antigens detected by different antibodies used at different RPPA platforms were, in general, almost as well correlated as those used at the same RPPA platform. These observations strongly suggest that the different instrumental set-ups and analytical workflows used at these representative RPPA platforms do not preclude the generation of comparable and reproducible data when using high-quality, validated antibodies.

Using different workflows, the layout of samples spotted on arrays differed among the three RPPA platforms, which prevented application of the same automated data normalisation method. Each research site therefore applied their own in-house data normalisation procedure tailored for the respective RPPA platform. All platforms normalised signal intensities for total amounts of printed protein, but the way they accounted for sample dilutions differed. The Paris platform applied non-parametric curve fitting to the serial dilutions of each sample, the Heidelberg platform fitted curves for control samples only and reported experimental samples to these curves and the Edinburgh platform computed a linear fit of the serially diluted samples. Despite these different approaches, we show that normalised data improves the distinction between concordant data (derived from antibodies recognising the same antigen class) and non-concordant data (derived from all antibodies regardless of target), as compared to raw data. Future efforts to develop normalisation pipelines that are compatible with multiple array layouts may further improve data reproducibility among platforms and expand opportunities for cross-platform validation of RPPA technology. Such cross-platform validation may have utility in the appraisal of robust markers of disease and therapeutic response and their application as prognostic or predictive biomarkers.

There is an outstanding need for the development of robust markers of response and resistance to targeted cancer therapy (Mueller et al. 2018, Giudice & Petsalaki 2019, Pierobon et al. 2019). Proteomic and phosphoproteomic datasets are well placed to complement genetics-based biomarker strategies by providing additional information on activation states of dynamic pathway signalling. However, the application of proteomic technologies presents a number of challenges, including consistent high-quality sample preparation, sensitive detection of low-abundance proteins and post-translational modifications, high sample throughput at reasonable cost and rapid turnaround time necessary for clinical application or drug discovery and development pipelines. The development of RPPA workflows attempts to address many of these challenges, yet the reproducibility of RPPA data generation and analysis across distinct RPPA platforms and research centres has not been extensively evaluated. We used an integrative RPPA approach to characterise a range of human breast cancer cell lines and their biochemical responses to two clinically relevant cancer drugs. The cell lines were chosen to represent different molecular subtypes of breast cancer, each of which having a different prognosis and treatment response (Sørlie et al. 2001). The sensitivity of these cell lines towards lapatinib and selumetinib and the expected changes in major signalling pathways are known (Konecny et al. 2006, Garon et al. 2010, O’Neill et al. 2012) and thus served as benchmarks in this study. We demonstrated that RPPA technology can identify expected protein markers for response to treatment and resistance in a robust and platform-independent manner. Understanding how resistance can be predicted and prevented is a major therapeutic challenge, and the use of proteomic approaches such as RPPA will, by defining the functional state of cells and tissues, enable the validation and assessment of resistance mechanisms in clinical samples (Creedon et al. 2014, Mueller et al., 2018).

High attrition rates in clinical drug development present significant challenges to drug developers, with only one in eight oncology drugs that enter clinical development in phase 1 achieving US Food and Drug Administration approval, and a 1-in-15 success rate when these candidate drugs are under evaluation in secondary oncology indications (Hay et al. 2014). The development of robust biomarkers is thus becoming an essential component of new clinical trial designs to guide patient selection, optimise dosing schedules and minimise ineffective or over-treatment. For many complex diseases, biomarkers at the genetic, proteomic, metabolomic and phenotypic levels are required to characterise individual patient disease and response to therapy sufficiently. Integration of multiple biomarker modalities with emerging computational and statistical approaches represents the future direction of personalised medicine strategies. However, the successful implementation of personalised medicine is dependent upon the validation and reproducibility of biomarker tests performed across distinct research centres and national boundaries.

Our data show that RPPA analyses of drug-treated breast cancer cells using distinct antibodies and different RPPA platforms can identify robust profiles of protein markers reporting signalling pathway responses to pharmacological inhibition. This suggests that the consistency of RPPA-based assays will enable the validation and assessment of treatment response and resistance mechanisms in clinical samples across international laboratories. These data provide, to our knowledge, the first extensive cross-platform validation of RPPA technology, which paves the way for further investigation and improvement of technology robustness.

## Methods

### Cell lines and cell lysis

MCF7, MDA-MB-231, MDA-MB-468, MDA-MB-453, HCC1954 and SKBR3 breast cancer cells were purchased from American Type Culture Collection and grown according to supplied instructions (Supplementary Table 1). For the preparation of cell lysates, cells were washed twice with ice-cold phosphate-buffered saline (PBS). Laemmli buffer (50 mM Tris-HCl (pH 6.8), 2% sodium dodecyl sulfate, 5% glycerol, 2 mM DTT, 2.5 mM EDTA, 2.5 mM EGTA, supplemented with 4 mM sodium orthovanadate, 20 mM sodium fluoride, Halt phosphatase inhibitor cocktail (Perbio) and cOmplete protease inhibitor cocktail (Roche)) was incubated at 100°C for 5 min and applied directly to cells. Samples were immediately incubated at 100°C for 10 min. Lysates were passed through a 25-gauge needle five times and clarified by centrifugation (18,000 × *g*, 10 min, room temperature). Clarified lysates were aliquoted and snap-frozen in liquid nitrogen prior to shipment to the various research sites. Protein concentration was determined using a reducing agent-compatible BCA kit (Pierce).

### Pharmacological inhibitor treatment

Lapatinib and selumetinib (Selleck Chemicals) were prepared as 10 mM stock solutions in DMSO. Cells were treated with 1 µM lapatinib, 1 µM selumetinib or DMSO in growth medium for 20 min or 24 h. Control cells were treated with DMSO in growth medium for 20 min.

### RPPA analysis

Samples were analysed at each research site using the respective in-house RPPA platforms as summarised in Supplementary Table 2. Biological triplicate lysates were serially diluted, if applicable, to produce a dilution series comprising four serial 2-fold dilutions of each sample (Supplementary Table 2). Sample dilution series were spotted onto Grace Bio-Labs ONCYTE nitrocellulose-coated slides (Sigma-Aldrich) in technical duplicate or triplicate under conditions of constant 70% humidity using an Aushon 2470 arrayer (Aushon Biosystems). Slides were hydrated in deionised water and blocked with blocking buffer (Supplementary Table 2). Slides were washed with Tris-buffered saline containing 0.1% Tween 20 (TBS-T) and incubated with validated primary antibodies diluted in blocking buffer at room temperature for 1 h or at 4°C overnight (Supplementary Table 3). Slides were washed with TBS-T and probed with secondary antibodies diluted in blocking buffer at room temperature for 30 min or 1 h. To amplify the signal, if applicable, slides were incubated with avidin, biotin and peroxidase blocking reagents (Dako) prior to primary antibody incubation and then with Bio-Rad Amplification Reagent (Bio-Rad) at room temperature for 15 min after secondary antibody incubation (Supplementary Table 2). Slides were washed with TBS-T. For staining of total protein with SYPRO Ruby, slides were washed once with 7% acetic acid, 10% methanol (15 min), twice with deionised water, once with SYPRO Ruby (Thermo Fisher Scientific) and once with deionised water. For staining of total protein with Fast Green FCF, slides were washed once with deionised water, once with 0.000005% Fast Green FCF (Sigma-Aldrich) and once with deionised water. Slides were allowed to dry at room temperature for 10 min prior to slide scanning. Slides were read using a GenePix 4000B (Molecular Devices), Odyssey (LI-COR Biosciences) or InnoScan 710-IR (Innopsys) scanner (Supplementary Table 2). The relative fluorescence intensity of each sample spot was quantified using MicroVigene (VigeneTech), GenePix Pro (Molecular Devices) or Mapix (Innopsys) software (Supplementary Table 2). Processed signal intensities were normalised using NormaCurve (Troncale et al. 2012), RPPanalyzer (Mannsperger et al. 2010) or spot-by-spot division of antibody signal intensity by total protein stain signal intensity (Supplementary Table 2).

### Dataset integration

Normalised RPPA data derived from each platform were binary-logarithm transformed, if necessary, and median centred antibody-wise. Unique antibody identifiers were assigned to each distinct antibody based on antibody reference, and antibody antigens were classified to account for recognition of up to two related protein or phosphoprotein isoforms or family members (e.g. antibodies targeting Erk1 and Erk1/2 were classified as recognising Erk1/2) (Supplementary Table 3). Data for antibodies targeting the same antigen class used on all three RPPA platforms were used for dataset integration; antigen classes represented by data derived from fewer than three RPPA platforms were excluded from further analysis.

### Unsupervised learning

Binary, agglomerative hierarchical cluster analyses of centred normalised abundances for proteins and phosphoproteins were performed using Cluster 3.0 (C Clustering Library, version 1.54) (de Hoon et al. 2004). Spearman rank correlation coefficients, Euclidean distances and Kendall tau coefficients were calculated and adapted as distances, if necessary. Distance matrices were calculated using pairwise average linkage. Hierarchical clustering results were visualised using Java TreeView (version 1.1.5r2) (Saldanha 2004). Principal component analyses were performed using Python (version 3.7.4) or Perseus (version 1.5.2.6) (Tyanova et al. 2016).

### Clustered antibody antigen mapping

For data-driven representation of antibody similarity across multiple RPPA platforms, we devised the clustered antibody antigen map. For all antibodies used in the integrative analysis, centred normalised abundances for proteins and phosphoproteins were clustered on the basis of Spearman rank correlation coefficient–based distance, Euclidean distance or Kendall tau coefficient–based distance using Cluster 3.0, computing distances with an average-linkage matrix. Clustered antibody (column) node memberships were stored for antibody mapping, and clustered antibodies were expanded in a second dimension according to antibody antigen classification. Antibodies targeting the same antigen class were indexed according to unique antibody identifier, enumerating from 1. A matrix of integer-indexed antibody identifier elements was generated according to clustered antibodies (columns) and corresponding antibody antigen classes (rows) (Supplementary Table 3). In this matrix, antibody (column) node memberships determined by clustering of RPPA data were preserved, and antibody antigen classes (rows) were ordered alphabetically. The resulting clustered antibody–antigen feature space that was used to annotate the integrated multi-platform RPPA data. Mapping results were visualised using Java TreeView.

### Statistical analyses

No statistical methods were used to pre-determine sample size. Spearman rank correlation coefficients were calculated for every pair-wise combination of antibody identifiers. Kernel density estimates were computed using R (version 3.4.1) (R Core Team 2017). Median Spearman rank correlation coefficients were compared using Fisher transformation and two-sided *z*-tests. Differentially regulated proteins and phosphoproteins were compared using two-tailed Student’s *t*-tests with artificial within-groups variance set to 1 and a permutation-based false discovery rate threshold of 5% (1,000 randomisations using Perseus).

## Acknowledgements

We thank Philippe Hupé for helpful scientific input, Patrick Poullet and Stéphane Liva for development and maintenance of data management tools at Institut Curie and Andrew H. Sims for discussions. A.B. was supported by Cancer Research UK. K.G.M. is supported by the Cancer Research UK Edinburgh Centre award (C157/A25140). The Institut Curie RPPA platform is supported by Cancéropôle Ile-de-France.

## Author Contributions

A.B., N.O.C., U.K., B.S. and L.d.K. conceived the study and designed the experiments. S.B., B.O., A.C. and K.G.M. generated the data. A.B., S.B., K.G.M., V.S., B.S. and L.d.K. analysed and interpreted the data. A.B. designed and implemented data integration and prepared the figures. A.B., N.O.C. and L.d.K. wrote the manuscript. All authors critically reviewed and approved the manuscript.

## Additional Information

### Supplementary information

accompanies this paper

### Competing interests

The authors declare no competing interests.

