## Supplementary Figures for "Integrative analysis of multi-platform reverse-phase protein array data for the pharmacodynamic assessment of response to targeted therapies"

### Supplementary Information

Supplementary Figure 1

Supplementary Figure 2

Supplementary Figure 3

Supplementary Figure 4

Supplementary Figure 5

Supplementary Table 1

Supplementary Table 2

Supplementary Table 3

Supplementary Table 4

Supplementary Table 5

Supplementary Table 6

[1]Cancer Research UK Edinburgh Centre, Institute of Genetics and Molecular Medicine, University of Edinburgh, Edinburgh, United Kingdom. [2]Division of Molecular Genome Analysis, German Cancer Research Center (DKFZ), Heidelberg, Germany. [3]Department of Translational Research, Institut Curie, PSL Research University, Paris, France. [4]U900 INSERM, Institut Curie, PSL Research University, Paris, France. [5]Present address: Pfizer Pharma GmbH, Berlin, Germany. [6]Present address: Sederma, Le Perray-en-Yvelines, France. [7]Present address: U1236 INSERM, Faculté de Médecine, Université de Rennes 1, Rennes, France. [8]Present address: NanoString Technologies, Inc., Seattle, Washington, United States of America. Correspondence and requests for materials should be addressed to A.B. or L.d.K.

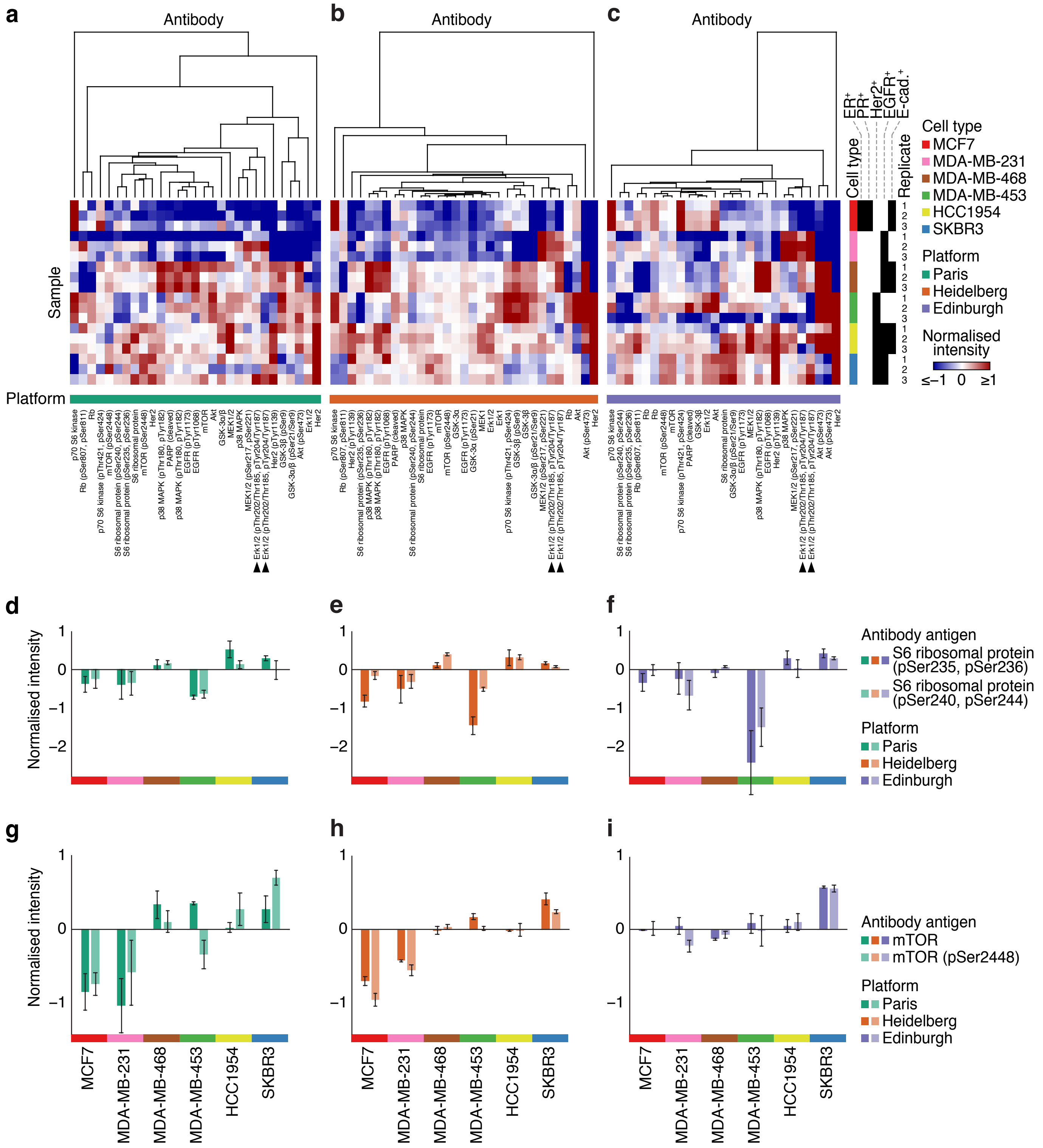


**Supplementary Figure 1.** RPPA data for distinct antibodies that recognise related antigens. (**a**–**c**) RPPA data derived from the Paris (**a**), Heidelberg (**b**) and Edinburgh (**c**) platforms were clustered on the basis of antibody-wise dissimilarity (Euclidean distance). Sample order and annotations are as for Fig. 1. (**d**–**f**) RPPA data for phosphorylated S6 ribosomal protein (dark bars, pSer235, pSer236; light bars, pSer240, pSer244) derived from the Paris (**d**), Heidelberg (**e**) and Edinburgh (**f**) RPPA platforms. The Paris and Edinburgh platforms used the same antibody that recognises pSer235, pSer236 (antibody identifier S6 ribosomal protein_pSer235,pSer236_a); the Heidelberg platform used the antibody with antibody identifier S6 ribosomal protein_pSer235,pSer236_b. All three platforms used the same antibody that recognises pSer240, pSer244 (antibody identifier S6 ribosomal protein_pSer240,pSer244). (**g**–**i**) RPPA data for mTOR (dark bars) and phosphorylated mTOR (light bars, pSer2448) derived from the Paris (**g**), Heidelberg (**h**) and Edinburgh (**i**) RPPA platforms. All three platforms used different antibodies that recognise mTOR (antibody identifiers for Paris, mTOR_b; Heidelberg, mTOR_c; Edinburgh, mTOR_a). Data derived from all three platforms for the same antibody that recognises phosphorylated mTOR (pSer2448) are shown (antibody identifier mTOR_pSer2448_a). Annotations are as for Fig. 1. Data are means ± s.e.m. ($n = 3$ independent samples). For further details, see Supplementary Table 4.

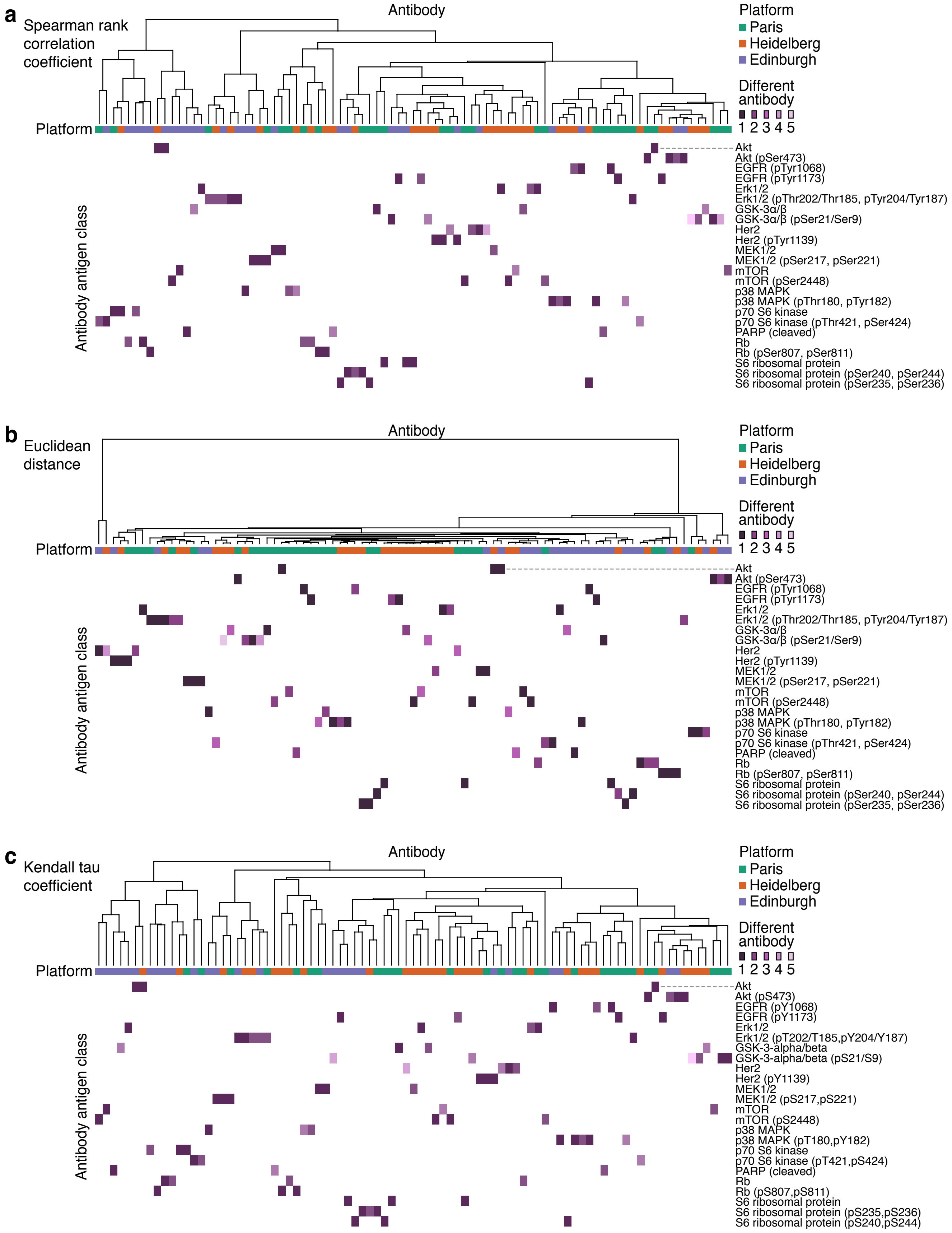


**Supplementary Figure 2.** Clustered antibody antigen maps of integrated RPPA data using different distance functions. (**a**–**c**) Annotated clustered antibody antigen maps of integrated multi-platform RPPA data of six breast cancer cell lines cultured under control conditions. The dendrograms representing antibody-wise dissimilarity are shown annotated with respective antibody antigen classes, and distinct antibodies that recognise the same antigen class (unique antibody identifiers) are indicated by different shades of purple, where applicable. Antibodies were clustered on the basis of Spearman rank correlation coefficient–based distance (**a**), Euclidean distance (**b**) or Kendall tau coefficient–based distance (**c**). Clustered antibody antigen map in **a** is the annotated version of that in Fig. 2. Antibody antigen classes are ordered alphabetically for clarity. Annotation bars indicate RPPA platform. For further details, see Supplementary Table 5.

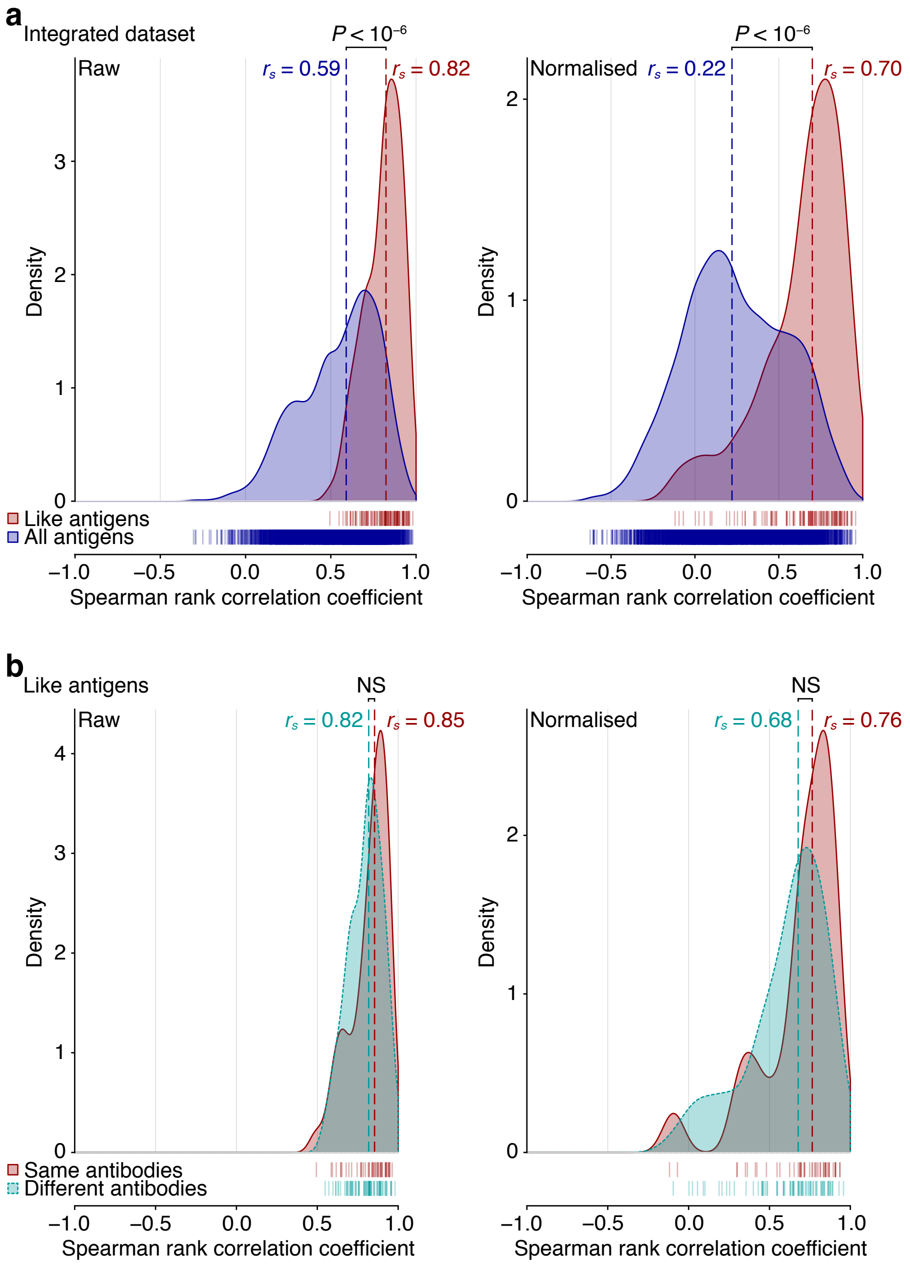


**Supplementary Figure 3.** Correlations of raw and normalised RPPA data used for integrative analysis of drug-treated breast cancer cell lines. (**a**,**b**) Kernel density estimates of Spearman rank correlation coefficients for every pair-wise combination of unique antibody identifiers were computed. Correlations of RPPA data derived from antibodies recognising the same antigen class (like antigens) were compared to those derived from all antibodies used in the integrated multi-platform RPPA dataset (**a**); for like antigens, correlations of RPPA data derived from the same antibodies were compared to those derived from different antibodies (**b**). Correlations of raw data (left panels) and normalised data (right panels) are shown. Spearman rank correlation coefficient data points for each set of comparisons are indicated by rug plots. For each set of comparisons, the median Spearman rank correlation coefficient ($r_s$) is shown (dashed lines). Differences in $r_s$ were assessed using Fisher transformation and two-sided $z$-tests. NS, not significant.

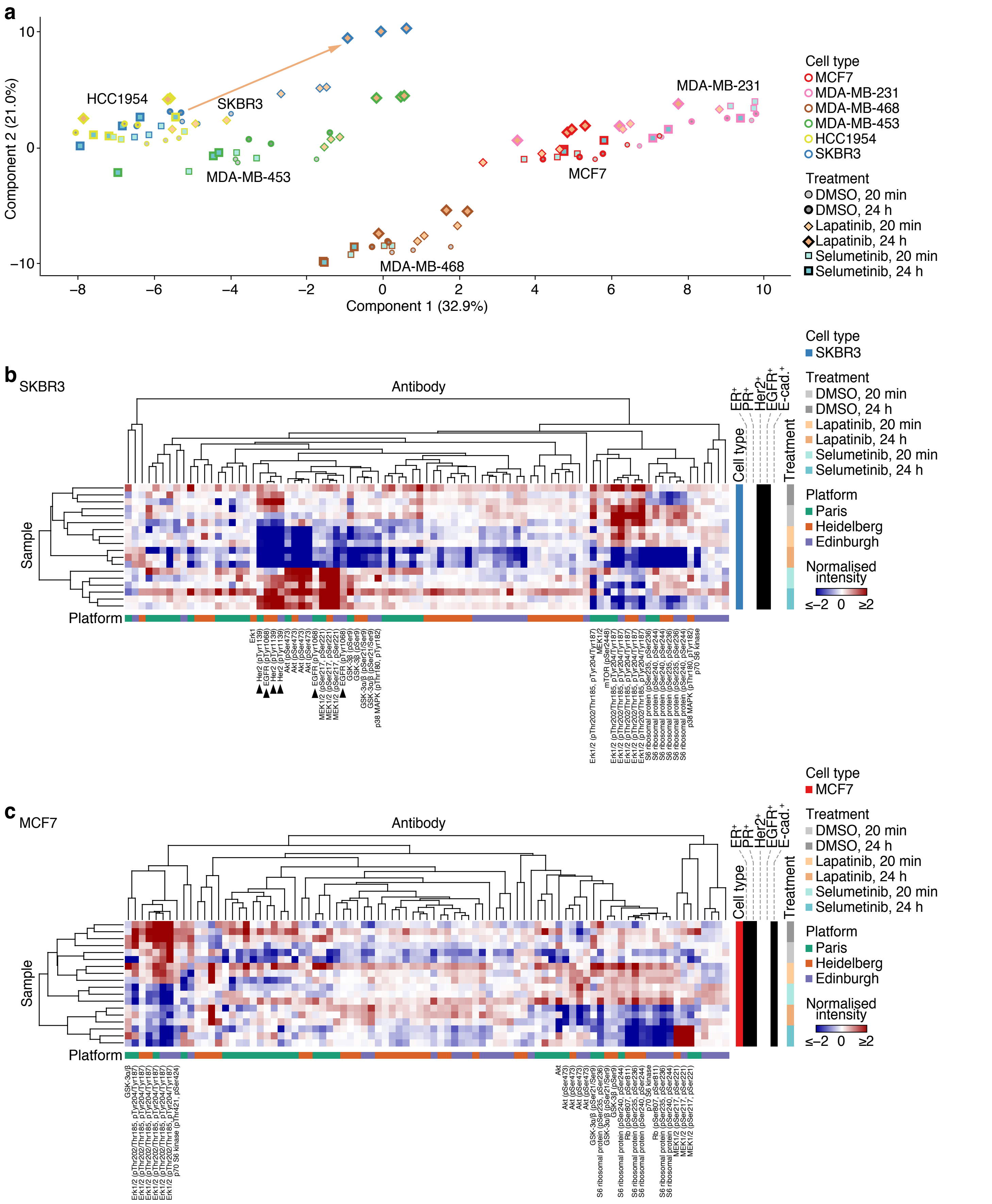


**Supplementary Figure 4.** Comparative analysis of drug-treated breast cancer cell lines. (**a**) Principal component analysis of integrated RPPA data of six breast cancer cell lines treated with lapatinib, selumetinib or vehicle control (DMSO) at two timepoints. (**b,c**) Integrated multi-platform RPPA data of drug-treated SKBR3 (**b**) and MCF7 (**c**) cells were clustered on the basis of antibody-wise and sample-wise dissimilarities (Spearman rank correlation coefficient–based distance). Annotation bars indicate cell type, cell type receptor status, drug treatment and RPPA platform. Clusters of antigens differentially regulated upon drug treatment are labelled; arrowheads indicate different phosphorylated Her2 and EGFR antibodies in the selected clusters.

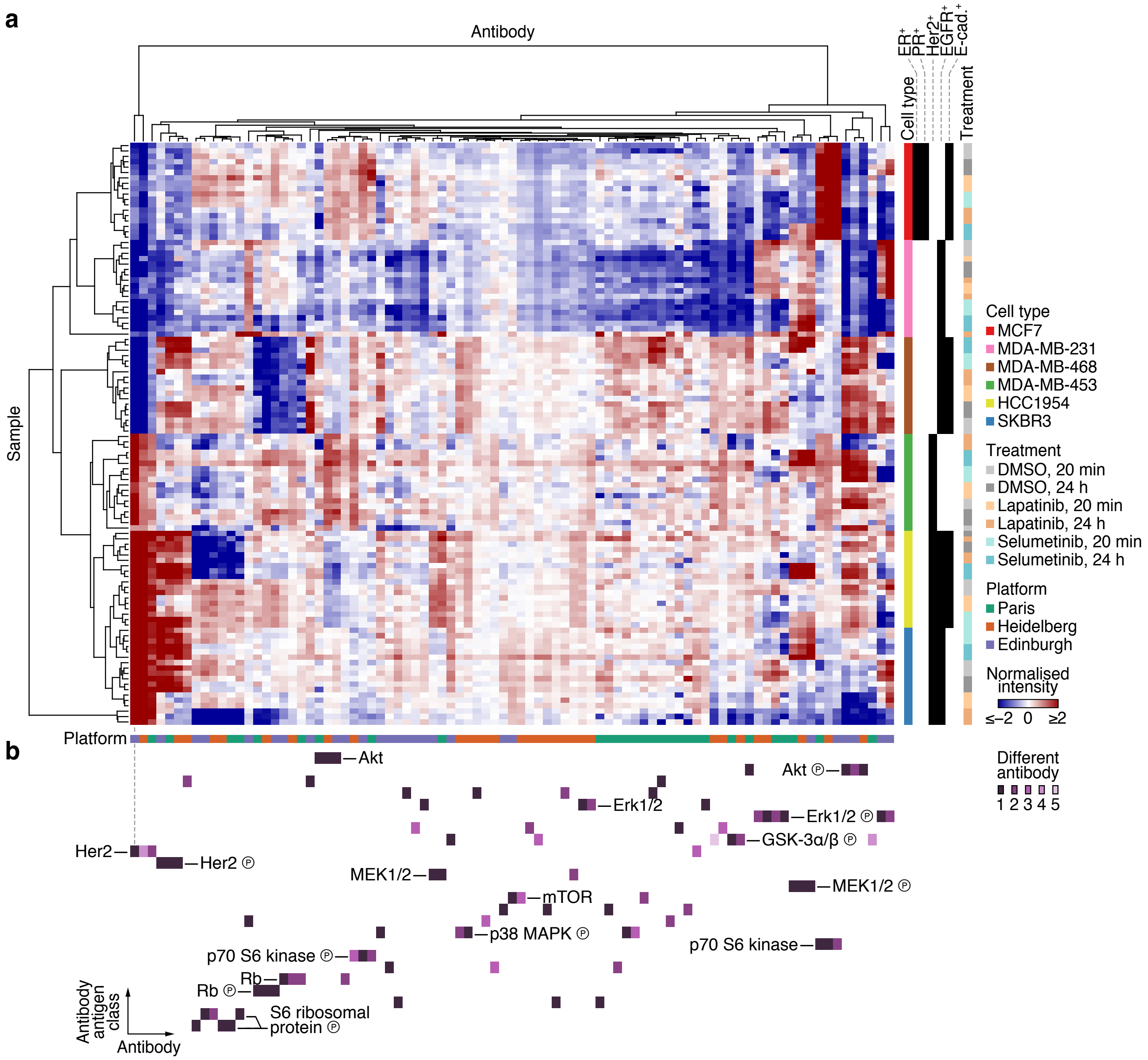


**Supplementary Figure 5.** Integrative analysis of drug-treated breast cancer cell lines. (**a**) Integrated RPPA data of six breast cancer cell lines treated with lapatinib, selumetinib or vehicle control (DMSO) at two timepoints. Data were clustered on the basis of antibody-wise and sample-wise dissimilarities (Euclidean distance). Annotations are as for Fig. 4. (**b**) Clustered antibody antigen map for hierarchical clustering results in **a**. Antibody antigen classes are ordered alphabetically for clarity. For further details, see Supplementary Table 6.
